# PIPP: Improving peptide identity propagation using neural networks

**DOI:** 10.1101/2021.12.05.471338

**Authors:** Soroor Hediyeh-zadeh, Jarryd Martin, Melissa J. Davis, Andrew I. Webb

## Abstract

Peptide identity propagation (PIP) can substantially reduce missing values in label-free mass spectrometry quantification by transferring peptides identified by tandem mass (MS/MS) spectra in one run to experimentally related runs where the peptides are not identified by MS/MS. The existing frameworks for matching identifications between runs perform peak tracing and propagation based on similarity of precursor features using only a limited number of dimensions available in MS1 data. These approaches do not produce accompanying confidence estimates and hence cannot filter probable false positive identity transfers. We introduce an embedding based PIP that uses a higher dimensional representation of MS1 measurements that is optimized to capture peptide identities using deep neural networks. We developed a propagation framework that works entirely on MaxQuant results. Current PIP workflows typically perform propagation mainly using two feature dimensions, and rely on deterministic tolerances for identification transfer. Our framework overcomes both these limitations while additionally assigning probabilities to each transferred identity. The proposed embedding approach enables quantification of the empirical false discovery rate (FDR) for peptide identification, while also increasing depth of coverage through coembedding the runs from the experiment with experimental libraries. In published datasets with technical and biological variability, we demonstrate that our method reduces missing values in MaxQuant results, maintains high quantification precision and accuracy, and low false transfer rate.

## Introduction

Liquid-chromatography-coupled tandem mass spectrometry (LC-MS/MS) is the leading technology for quantitative analysis of proteins expressed in complex biological samples. Proteins in cell or tissue lysates are first prepared for analysis by extracting the protein content, followed by enzymatic digestion, converting them into peptides. Peptides are separated using liquid chromatography which is interfaced with the source of the mass spectrometer, where they are ionised and converted to gas phase. The separated and ionised peptide precursors are subjected to mass analysis in a mass spectrometer. During conventional data-dependent acquisition (DDA), peptide ions are sampled for fragmentation and identified from the spectra produced by tandem mass (MS/MS) analysis using peptide identification software (Cox et al., 2011). More recently, high resolution Ion Mobility (IM) has been incorporated in the instrument platform and providing an additional dimension for peptide separation. The mass, charge, retention time and IM collectively can define the identity of peptides existing in a sample. It is known that more than 100,000 detectable peptide species elute in single shotgun proteomics runs (Michalski et al., 2011). In DDA, however, the mass spectrometer only selects a small subset (usually 10) of the most abundant peptides for sequencing by MS/MS at each MS1 survey scan in a run. This compromises consistent identification of peptides across runs, as the sets of peptide precursor ions selected for sequencing could be different between runs. The low sampling efficiency and stochastic nature of intensity-dependent sampling of peptide ions for MS/MS analysis limits depth of protein coverage and hinders quantification of low abundance ions in complex samples, leading to the prevalent problem of missing values.

To alleviate the missing values problem in mass spectrometry data, peptide identifications are transferred between runs to increase the number of peptides and proteins quantified across runs. This is known as Peptide Identity propagation (PIP) (Zhang et al., 2016; 2017), or match-between-runs (MBR) (Cox et al., 2014). When a peptide is sequenced in a run, its MS1 attributes such as *m/z*, retention time and Collision Cross Section (CCS) are available to the experimenter. Data completeness can be improved by transferring peptide identifications from runs where the peptide is identified by sequencing (MS/MS) to runs where it is not sequenced or identified, but is detected at the MS1 level based on the similarity of the MS1 attributes. Existing *identification transfer* approaches transfer sequence information to a run if a peptide-like signal (precursor feature) is present for which the MS1 measurements, that is *m/z* (denoting mass over charge), retention time and IM, are within a predefined tolerance of MS1 attributes of a peptide identified by sequencing (Zhang et al., 2016; Yu et al., 2020a; Shen et al., 2018; Yu et al., 2020b). Peak tracing and sequence propagation are typically carried out in two dimensions: Yu et al. (2021) transfer sequence information by peak tracing in *m/z* - RT dimensions, while Demichev et al. (2021) propagate precursor identities by peak tracing in *m/z* - CCS dimensions. Some approaches such as Zhang et al. (2016) transfer identifications from the run with the largest number of identified peptide sequences, whereas Yu et al. (2020a) transfer identifications from the top *k* correlated runs. Therefore, the accuracy of the existing approaches is limited by the selection of runs used as a *reference* for PIP and by the specified tolerance. These approaches are limited by the lack of any probability measure that could serve as a confidence estimate by which likely false positives could be filtered out. Recently, May et al. (2018) used deep neural networks and learned an embedding of millions of mass spectra. The learned embedding has the potential utility to transfer identifications among nearby spectra in the embedding. However, statistical confidence estimation procedures for peptide identity propagation remain to be developed for this approach.

We propose a new approach for identification transfer using neural networks that involves embedding MS1 attributes of peptide peaks into a higher dimensional space. This higher dimensional space (embedding) is learned through a classification framework, where the classes are peptide identifications. Since the embedding is learned in a fully-supervised classification framework, the embedded representations of precursor features are optimized to capture attributes that define the identity of a peptide on the basis of its MS1 measurements.

Here we present *peptideprotonet*, a variation of prototypical networks (Snell et al., 2017) trained in a few-shot learning framework that learns embedded representations of peptides based on the MS1 measurements. We refer to the peptides not sequenced by MS/MS, but detected at the MS1 level as *query precursor features* hereafter. The query precursor features detected by a peptide signal detection algorithm, such as MaxQuant, are mapped to an embedding space learned by the model. The sequence and charge information are then propagated from identified (sequenced) peptides to query precursor features based on the similarity of their representations in the embedding space. The embedding approach for identification transfer obviates the reliance of PIP on deterministic tolerance thresholds and choices of reference runs, performs propagation in more than two or three dimensions and assigns confidence scores to each transferred identification. In addition, the proposed identification transfer approach, Peptide Identity Propagation with Protonets (*PIPP*), allows to co-embed multiple runs and peptide identification libraries to increase depth of coverage and estimate false-discovery rate (FDR) in identification transfers.

We demonstrate applications of the *peptideprotonet* model on precursor features reported by MaxQuant in LC-IMS-MS/MS (trapped ion mobility mass spectrometry) data. In contrast to May et al. (2018), our embedding model and PIP framework does not require access to raw mass spectra, assigns statistical confidence measures to individual identification transfers and enables computation of the empirical FDR. Since MaxQuant is currently the only available software that reports all precursor features that it has detected regardless of the MS/MS outcome of the precursor feature, and there is currently no other comparable embedding based PIP approaches, we have compared our results with MaxQuant MBR algorithm in a number of DDA datasets acquired by Parallel Accumulation - Serial Fragmentation (PASEF) with biological and technical variability, using published MaxQuant results.

## Results

### Peptideprotonet: an embedding approach to peptide identity propagation

Peptideprotonet is a new approach for MS1-based peptide identity propagation between runs of an experiment that transfers identifications between precursor feature signals based on the similarity of MS1 attributes in a higher dimensional state space, rather than the 2- or 3-dimensional (*m/z* - RT, *m/z* - CCS, or *mz*-RT-CCS) search spaces (Prianichnikov et al., 2020; Demichev et al., 2021; Yu et al., 2021) in which existing methods measure similarity. This is achieved by embedding the MS1 attributes of the precursor features onto larger dimensions using Neural Networks (Methods and Materials). The embedding of a precursor feature is a new, higher-dimensional representation (vector) of its MS1 measurements, and accommodates richer information on the similarity of precursor features in MS1.

Peptideprotonet is based on the idea that more than 100,000 peptides elute in a single LC-MS run, but the majority are not accessible for fragmentation. The precursor features are assumed to have been detected by a feature detector. Our approach takes precursor features that are detected by MaxQuant in MS1, but are missing the MS2 information, and defines the probability that any such precursor feature could be assigned to peptides with non-missing MS2 information - that is peptides identified by fragmentation within the experiment. We trained a deep neural network model (Methods and Materials) that aligns retention times for peptides quantified in two large DDA-PASEF datasets of whole-proteome digests from HeLa, Yeast, Ecoli, CElegans and Drosophila (Figure 1a). This alignment (Figure 1b, middle panel) is achieved by learning an embedding of precursor features in which peptides from the same precursor (sequence-modification-charge) will have similar embedding vectors (Figure 1c, and Methods and Materials). The model was learned using MaxQuant results (the *evidence* tables) and is designed to improve confident identifications between runs using only the MaxQuant results. On a number of published datasets with biological and technical variability acquired with different retention gradient lengths, we demonstrate that our embedding-based PIP framework improves data completeness in the MaxQuant results, while maintaining high quantification precision and accuracy, and low false discovery rates (FDR).

**Figure 1.**
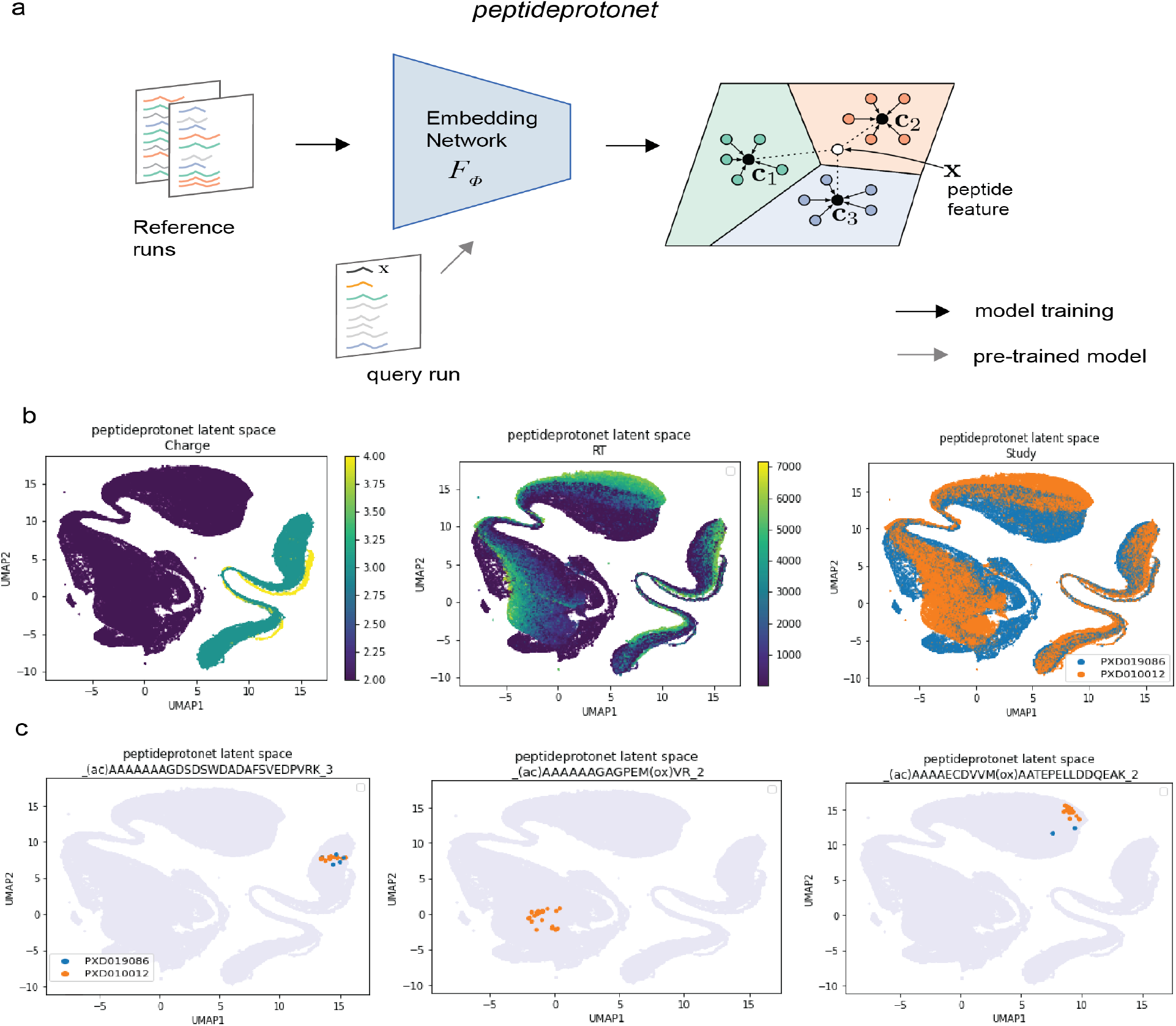
The peptideprotonet model. a) Schematic illustrating the peptideprotonet model. Deep Neural Networks are used to learn new representations of TIMS-MS1 measurements that align peptide precursor sequences identified by fragmentation in two comprehensive studies. Through such alignment, the network learns a model of each peptide. The model can be used to determine the identity of a precursor feature that is detected in MS1 by not sequenced in PASEF, therefore enhancing match-between-run and improving data completeness. The new representations are known as embedding. b) The embedding space of peptideprotonet colored by precursor charge, retention time and the study of origin. c) The embedded representations of three peptide precursors are highlighted. The proximity of embedded representation for peptide precursors identified in both studies indicate the goodness of alignment, hence the model.

The pre-trained model is applied to precursor features reported in the *allPeptides.txt* results of MaxQuant (MBR disabled) to obtain the embedded representation of precursor features in a query study. The identifications are then propagated from features identified by MS2 (reported in the *evidence.txt* results) to precursor features for which MS2 information is not available. This propagation is based on the similarity of their embedding vectors. A confidence score (probability) is assigned to each propagation. This entails that less confident propagations can be identified and discarded from the results. Therefore, unlike existing approaches, our approach does not rely on arbitrary thresholds such as ppm and/or RT tolerance between donor and receiver precursor features, as the propagation is probabilistic.

The nature of the model allows co-embedding of the query runs (runs from the experiment) with one or more experimental libraries of peptide identifications, for example, *evidence.txt* results from a previous study. In the performance evaluation results that follow, we co-embed runs from query (Human or HeLa) studies with HeLa and Yeast libraries and perform peptide identity propagation in two passes. In the first pass, propagations with confidence scores greater than a pre-specified threshold, *thr*, are assigned to within-experiment peptides. If the confidence score for a precursor feature is below the threshold, the feature is compared against peptide identifications from the co-embedded libraries in the second pass. Any new HeLa identifications transferred to query study would increase depth of coverage, if the transferred peptides belong to proteins that were not originally quantified in the experiment. Furthermore, any Yeast identifications transferred to HeLa/Human runs are false positives, and can be used to measure identification FDR empirically, that is

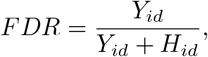

where, *Y*_*id*_ is the number of Yeast identifications (ids) and *H*_*id*_ is the number of Human peptide ids. A similar strategy has been recommended in Data Independent Acquisition (DIA), where the reference libraries are augmented with identifications from other species such as Arabidopsis (Demichev et al., 2021). The number of Arabidopsis identifications (calls) detected in runs are used to quantify the empirical FDR for identifications. This approach replaces the conventional target-decoy approaches (Nesvizhskii, 2010) for FDR estimation in DDA.

### Improved data completeness at peptide level

We applied peptideprotonet+PIP to four HeLa cell lysate replicates searched and quantified by MaxQuant with MBR disabled (MaxQuant MBR-) (Figure 2a). We used MaxQuant results published by Meier et al. (2018). MaxQuant results reported 53639 identifications in total. We defined data completeness as the proportion of total identifications with no missing values in all replicates. We observed that data completeness increased from 44% to up to 83% compared to MaxQuant MBR-, while there was up to 2.3% increase in peptide CV and up to 0.8% increase in protein CV (peptide CV for MaxQuant MBR-results was 1.2%, protein CV for MaxQuant MBR-was 2.1%, max peptide CV in peptideprotonet+PIP results was 3.5% corresponding to thr=0, and max protein CV in peptideprotonet+PIP was 2.9%). Overall, we observed 39% increase in data completeness, while peptide and protein CVs remained below 5%, demonstrating high peptide and protein quantification precision in PIP results.

**Figure 2.**
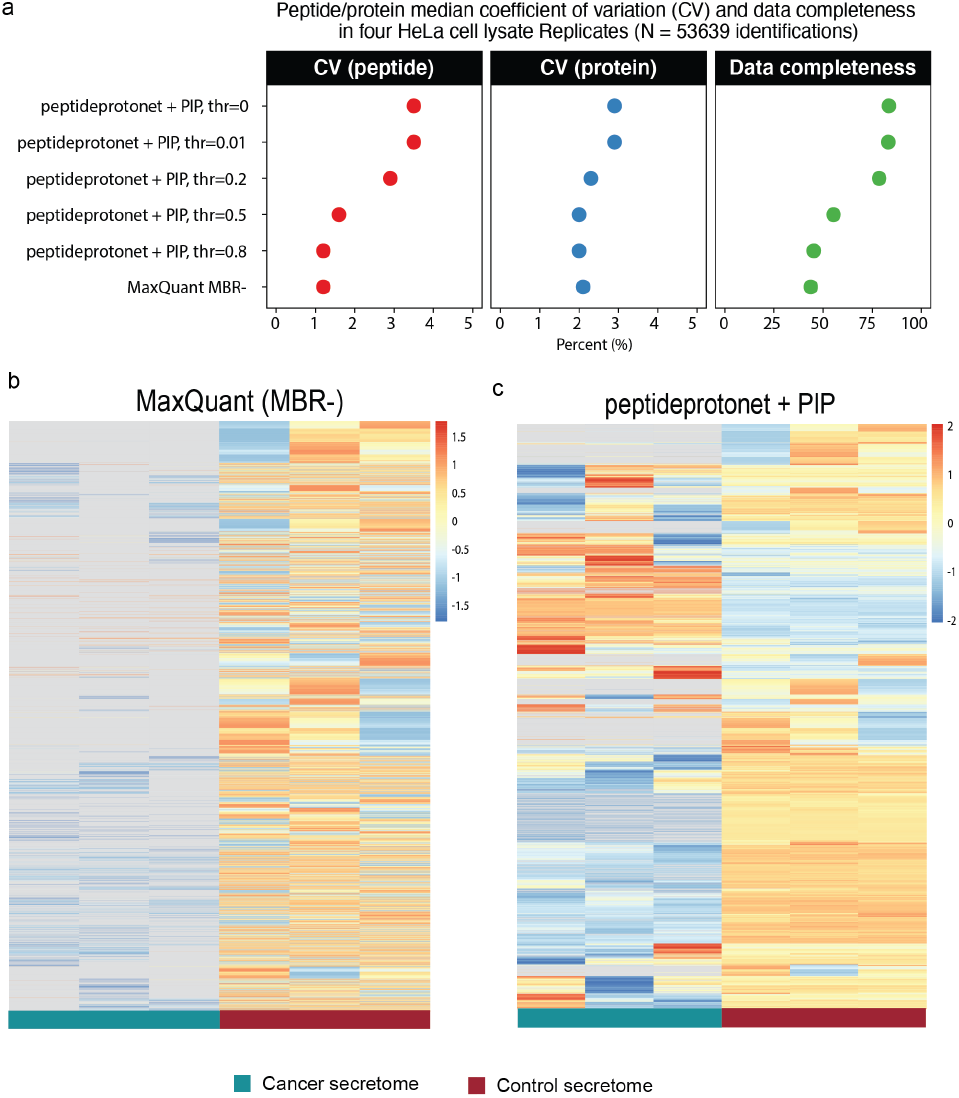
Peptideprotonet + PIP enhances data completeness in homogeneous technical runs and a heterogeneous dataset with biological variability. a) peptide and protein coefficient of variation (CV) and data completeness in four HeLa cell lysate replicates for different confidence score thresholds are compared to MaxQuant without Match Between Run (MaxQaunt MBR-). We used MaxQuant results published by Meier et al. (2018). Note evaluations are on the same number of identifications. Lower CVs are desired. b) Heatmap of scaled peptide log-intensity for 7728 peptides quantified completely in control OSCC secretome that are partially or completely missing in cancer OSCC runs. Gray areas represent missing values. Less gray areas are desired.

We assessed data completeness in a more challenging dataset with biological variability and small number of replicates. We identified an oral squamous cell carcinoma (OSCC) dataset studying T cell populations in tumor microenvironment. This dataset contains three secretome proteomes of OSCC tumor microenvironment, and three secretome proteomes of non-malignant samples. MaxQuant MBR-identified 30843 peptides. We defined data completeness as described earlier. Data completeness was increased from 29% to 78% when peptideprotonet+PIP (*k*=10, *thr* = 0) was applied to MaxQuant MBR-results. We identified a set of 7728 peptides with a measurement in all three control samples, that were missing partially or completely in cancer samples (Figure 2b). The set was selected to assess the performance of the method when missing values are likely not at random (MNAR). We were able to assign an intensity value to 52% of these peptides that with likely MNAR missingness that were not originally quantified in MaxQuant MBR-results.

### Increased depth of coverage and low false transfer rates

In addition to match between runs by means of within-experiment peptide identity propagation, peptideprotonet+PIP can be used to increase depth of protein coverage, and identify proteins that were not originally detected in MaxQuant results. This is achieved by co-embedding the query data (dataset of interest) with an experimental library; for example, a library of identifications, that is MaxQuant results, from a previous study or experiment.

Lim et al. (2019) evaluated false transfers in MaxQuant match between run algorithm by designing a two-organism DDA dataset. This dataset contains 20 replicates from a mixture of Human (90%) and Yeast (10%) proteins and 20 replicates of Human-only proteins. They evaluated false transfers by Yeast identifications transferred to Human runs by match between run. We evaluated false transfer rate in the presence of biological variability (the T cells OSCC dataset) and in the absence of biological variability (the ten HeLa cell lysate replicates) by co-embedding each of the two datasets with a Yeast library, and quantifying the proportion of Yeast identifications propagated to Human runs at various confidence thresholds and for different values of *k* (Table 1). We used MaxQuant (MBR enabled) results published by Prianichnikov et al. (2020) for the ten HeLa replicates dataset. In this case, the library is obtained by (randomly) down-sampling the Yeast identifications used at the training step. We also defined coverage per run as the number of identifications in the run over all identified peptides. We observed that the proportion of Yeast identifications transferred to the runs was generally below 5% for all possible combinations of *k* and *thr* in both datasets, and was higher for the less strict confidence thresholds such as *thr*=0.01, that is propagations with larger than 1% confidence threshold. Interestingly, we observed that at confidence score of 0.8 (i.e propagations that are more than 80% confident), there were no Yeast identifications transferred to the query runs, demonstrating that the peptideprotonet+PIP is a highly accurate framework for reducing the burden of missing values, and highlighting the importance of confidence measures in these data.

**Table 1.** False transfer rate and peptide coverage per run False transfer rate, mean coverage, median total identifications and new identifications found in a run after PIP in the absence of biological variation (HeLa cell lysate replicates), and when biological variation is present (T cells OSCC dataset). Each of the two datasets were co-embedded with a Yeast and HeLa library. False transfer rate is the proportion of Yeast identifications in HeLa/Human runs after PIP. Coverage is number of identifications in the run over all identified peptides. New identifications are HeLa identifications transferred from the HeLa library that were not previously detected in the experiment. Median identification is median of total identifications per run after PIP. Lower false transfer rate and higher coverage and identifications are desired. MaxQuant results for the HeLa cell lysate replicates are published by Prianichnikov et al. 2020.

| Threshold | HeLa cell lysate replicates 2hr gradient (n = 10)* |  |  |  | T cells OSCC (n = 6)† |  |  |  |
| --- | --- | --- | --- | --- | --- | --- | --- | --- |
|  | False transfer rate | Mean Coverage | New identifications | Median identifications | False transfer rate | Mean coverage | New identifications | Median identifications |
| **k = 5** |  |  |  |  |  |  |  |  |
| 0.00 | 0% | 0.909 | 0 | 57634 | 0% | 0.807 | 0 | 24978 |
| 0.01 | 1.6% | 0.908 | 1982 | 59544 | 0.8% | 0.799 | 928 | 25696 |
| 0.05 | 1.6% | 0.902 | 2146 | 59416 | 1% | 0.786 | 1218 | 25582 |
| 0.20 | 0.9% | 0.875 | 1524 | 57128 | 0.6% | 0.735 | 1049 | 23828 |
| 0.50 | 0.2% | 0.739 | 484 | 47661 | 0.2% | 0.593 | 318 | 18681 |
| 0.80 | 0% | 0.679 | 26 | 43364 | 0% | 0.547 | 12 | 16928 |
| **k = 10** |  |  |  |  |  |  |  |  |
| 0.00 | 0% | 0.860 | 0 | 54571 | 0% | 0.756 | 0 | 23400 |
| 0.01 | 0.9% | 0.860 | 688 | 55282 | 0.3% | 0.755 | 198 | 23572 |
| 0.05 | 1.3% | 0.859 | 1162 | 55660 | 0.6% | 0.751 | 484 | 23743 |
| 0.20 | 1.1% | 0.847 | 1348 | 55116 | 0.7% | 0.728 | 810 | 23364 |
| 0.50 | 0.3% | 0.786 | 607 | 50674 | 0.3% | 0.641 | 425 | 20255 |
| 0.80 | 0% | 0.702 | 38 | 44848 | 0% | 0.556 | 24 | 17258 |

Note:
The figures are based on peptide identifications per run.
\* Mean peptide coverage reported by MaxQuant is 0.67 before PIP;
† Mean peptide coverage reported by MaxQuant is 0.54 before PIP

The two datasets were additionally co-embedded with a random sub-sample of HeLa identifications in the training data. The identifications transferred from the co-embedded HeLa library that were not previously detected in the experiment were defined as new identifications. As expected, more new identifications were transferred to the query runs at less restrict thresholds, except for *thr* = 0, where the framework only transfers within-experiment identifications and no new identifications are allowed. The maximum mean coverage was also observed at this threshold. Interestingly, transfer of new identifications did not deteriorate the coverage. For example, in the HeLa cell lysate results with *k*=5, there was a median of 2146 new identifications at *thr*=0.05 and mean coverage increased from 67% in MaxQuant MBR-results to 90%. This demonstrates that the addition of new identifications did not increase the proportion of missing values.

The median identifications in MaxQuant MBR+ results of the ten HeLa cell lysate replicates was 42742 identifications. Peptideprotonet+PIP increased median identification by 10666 ± 5502 identifications. The Tcells OSCC dataset was quantified with MaxQuant MBR-, which resulted in median identifications of 16770 peptides. We observed an increase of 5504 ± 3154 identifications in peptideprotonet+PIP results.

When *thr*=0, every precursor feature is associated with a peptide sequence from the experiment irrespective of confidence score. Therefore, no new identifications are acquired from co-embedded libraries, and false transfer rate and number of new identifications are consequently zero. In general there are less transfers as *k* increases. This is because the standard deviation of distances (equation 1) increase as the search neighbourhood is expanded, confidence probabilities become smaller and it becomes more difficult to reach a confidence threshold. Further, smaller values of *k* encourage local propagations, while larger values of *k* encourage nonlocal propagations (that is transfers from prototypes that are further away from the embedded representation of precursor feature). This justifies higher mean coverage and median identifications for *k*=5 results in Table 1, as small *k* encourages local transfers. For *k*=10 we observed a higher number of new identifications with more than 80% confidence (i.e. *thr*=0.8) compared to *k*=5, confirming that propagation tend to be from a larger neighbourhood (non-local) as *k* increases.

### Quantification precision comparison using ten HeLa cell lysate replicates

We used the ten HeLa cell lysate replicates to compare peptide intensity precision after PIP to MaxQuant MBR+ results (Table 2). We observed up to 2% increase in peptide coefficient of variation (CV) in evaluations, while peptide data completeness increased up to 84% compared to 35.5% in MaxQuant MBR+ results. We also extended the evaluations to protein-level quantification. We defined quantified proteins as proteins with non-zero and non-missing values in at least two runs. Protein abundances were calculated from peptide intensities using top-N approach. We could quantify on average 763 ± 616 more proteins with peptideprotonet+PIP compared to MaxQuant (Table 2). For any choice of *k*, total peptides, quantified proteins and data completeness increased as the confidence score threshold was relaxed. This was, however, associated with increased coefficient of variation. This could be explained by identifications being generally intensity-dependent, where CV generally inversely correlates with intensity (Liu et al., 2015; Al Shweiki et al., 2017; Mahoney et al., 2011). The coefficient of variation, however, was consistently below 5%, concordant with PIP results in the four HeLa cell lysate dataset by Meier et al. (2018) (Figure 2a). This demonstrates that peptideprotonet+PIP framework does not compromise the precision of peptide quantifications, and can substantially improve data completeness. At *thr*=0, the number of total identifications is less than small thresholds such as 0.01 and 0.05 as no new identifications are acquired. This also explains why the largest proportion of complete data is observed at 0 threshold. On the other hand, at stringent thresholds such at *thr*=0.8, the number of total peptides is less than those for confidence scores between 0.01 and 0.5, as only highly confident propagations are retained in the results.

**Table 2.**
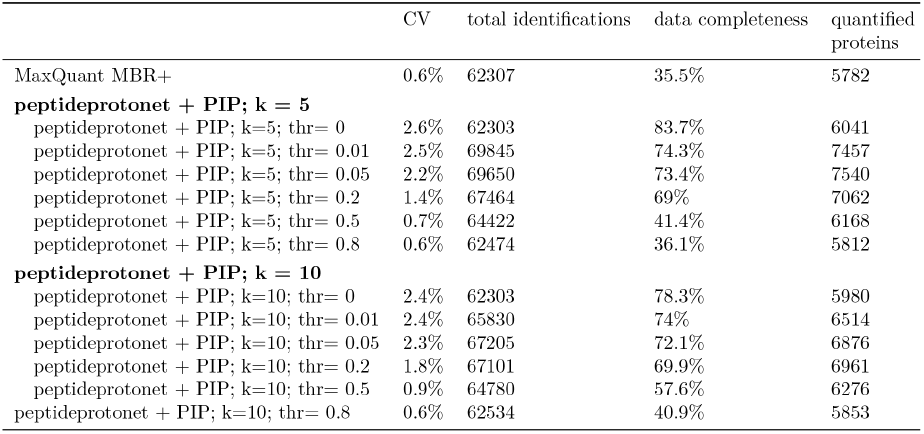
Peptide coverage, coefficient of variation (CV) and data completeness in ten HeLa cell lysate replicates Peptide coefficient of variation, number of identifications and data completeness, and quantified proteins in the ten HeLa cell lysate replicates. *Total identifications* is all the identified peptides in the experiment. *Data completeness* is the proportion of peptides with non-NA values in all the runs. *Quantified proteins* is the number of proteins with non-zero and non-NA values in at least two runs. Note that this table complements the coverage and false transfer rate results for the ten HeLa cell lysate dataset in Table 1. Lower CV values, larger number of identifications and quantified proteins, and higher fraction of complete data are desired.

### Quantification accuracy comparison using a three-organism hybrid proteome

We used a three-organism proteome to compare peptide fold-change estimation accuracy in peptideprotonet+PIP results with MaxQuant MBR+ results, when the ground-truth is known (Figure 3). The dataset contains six runs from two conditions A and B, where a mixture of Human, Yeast and E.coli proteins are spiked at known ratios. The ratios in B versus A are 1:1 for Human proteins, 4:1 for E.coli proteins and 1:2 for Yeast proteins. We applied peptideprotonet+PIP to MaxQuant results published by Prianichnikov et al. discarding any identifications that were not detected by MULTI-MSMS, which is equivalent to MBR-. We only retained propagations with confidence score greater than 0.5 (that is more than 50% confident) in the results. The comparison with MaxQuant MBR+ in the evaluations were done using the full published results by this study. For this dataset, we defined quantified proteins as proteins with non-zero, non-missing values in all six runs. Protein abundances were calculated from peptide intensities using top-N approach. We observed that our framework could quantify 412 more proteins than MaxQuant MBR+ at *k* = 5, and 1015 more proteins at *k* = 10 compared to MaxQuant. The peptide log_2_ fold-changes in each organism were centered around the expected ratio. This demonstrates that peptide identity propagation by peptideprotonet+PIP did not compromise the accuracy of peptide fold-changes. However, we did observe more outliers as *k* was increased.

**Figure 3.**
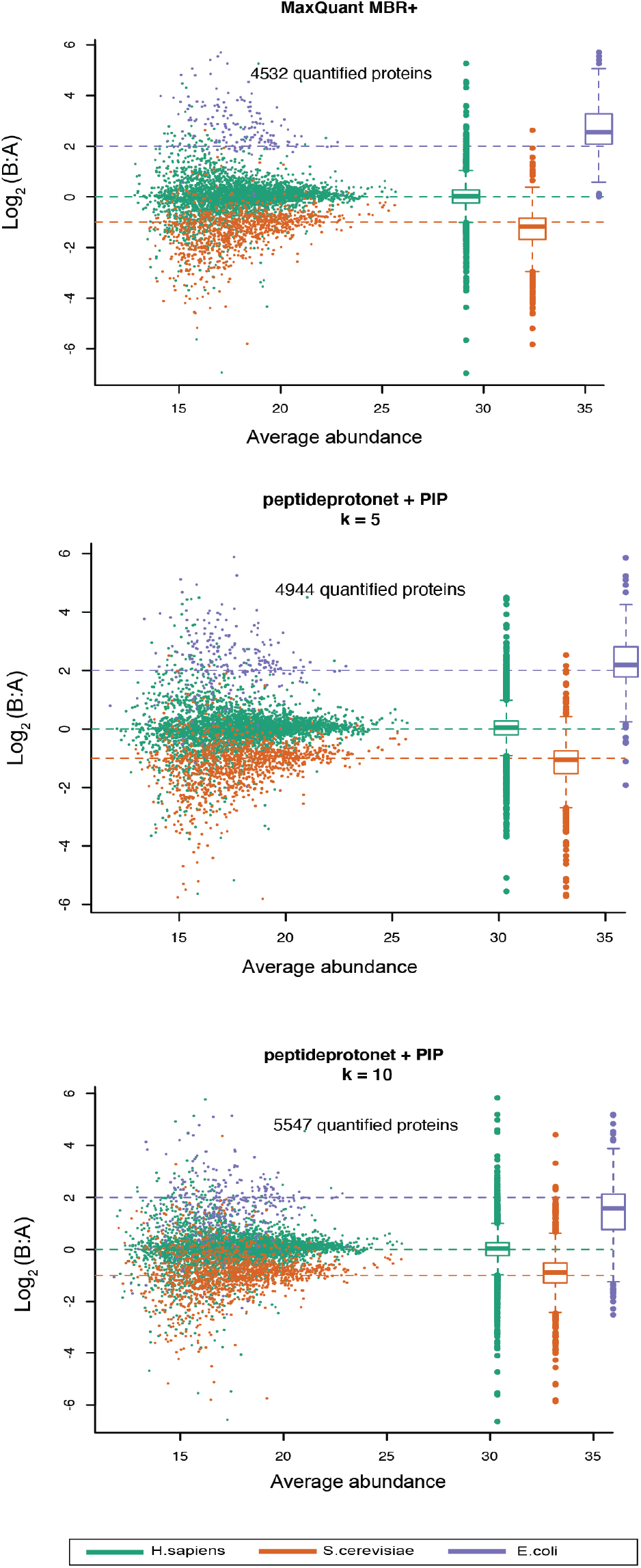
Protein quantification by MaxQuant and peptideprotonet+PIP in a hybrid Human, E.coli and Yeast proteome. The proteome ratios in condition B vs condition A are 1:2 (Yeast), 1:1 (Human) and 4:1 (E.coli). Box plots of protein log fold-changes are shown to the right of each scatter plot. MaxQuant results are published by Prianichnikov et al. 2020.

## Discussion

We introduced an embedding-based approached for identification transfer to match between runs using MS1 measurements. We developed a propagation framework for DDA-PASEF datasets that only requires MaxQuant results and does not require substantial reprocessing or access to raw spectral data. Using a number of published datasets with technical and biological heterogeneity acquired with different retention gradient lengths, we demonstrated that our embedding-based PIP framework improves data completeness in the MaxQuant results and maintains high quantification precision and accuracy.

Our framework overcomes the limitations of existing workflows by performing local (small *k*) and global (large *k*) propagation in more than two feature dimensions, dispensing with deterministic tolerances, and assigning probabilities to each transferred identity. In addition, the confidence scores from PIP results can be used as weights into *limma* linear models to account for uncertainty in the assignment of identifications, and improve power and reliability of differential abundance analysis. We were unable to evaluate the performance of this weighting strategy in the present work, however, as published controlled mixture PASEF datasets are currently lacking.

Our embedding approach enables quantification of the empirical FDR for peptide identification and increase in depth of coverage through co-embedding the runs from the experiment with experimental libraries. This flexibility in co-embedding multiple runs and libraries makes this approach particularly applicable to dia-PASEF data, and lays the foundation for future developments in embedding-based peptide identification.

## Materials and Methods

### The learning framework for peptideprotonet model

When proteins are digested into peptides in the sample preparation step, depending on the type of enzymes used and the pH of the digestion solution, the enzyme can cut the protein at different sites. Therefore, the generated peptide sequences can vary from run to run. Indeed, the occurrence of the same peptide sequence (i.e. the number of runs the peptide sequence is quantified in) with the same modification and charge state can be as low as one. In addition, retention time gradient length can vary between studies. The retention time of a peptide is a function of gradient length; that is the same peptide elutes at different times in 30 minute (short) gradient length experiments, compared to 120 minute (long) gradient length experiments. Due to the small and highly imbalanced number of data points available per identification (peptide sequence) for training, the model has to learn from only a few peptide occurrences. This is known as few-shot learning. We chose Prototypical Networks (Snell et al., 2017) approach in order to learn embedded representations of MS1 attributes, as they have proven to be highly successful in few-shot learning classification tasks.

### Prototypical Networks

Prototypical Networks or Protonets, learn an embedding function *f*_*ϕ*_ : ℝ^*D*^ → ℝ^*M*^ with learnable parameters *ϕ*, which maps the data points (precursor features in MS1) from the D-dimensional input feature space into an M-dimensional embedding feature space, with M *>* D. During training, a support set of labelled data points from each class are presented to the model, which maps them into the embedding space. A prototype embedding vector *c*_*k*_ is then computed for each class *k*, by averaging the embedded vector representations of the points in the support set for class *k*:

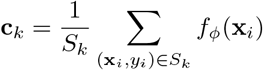

where *S*_*k*_ = {(**x**_1_, *y*_1_), …, (**x**_*N*_, *y*_*N*_)} is a small set of *N*_*k*_ labeled examples from class *k* where each *x*_*i*_ ∈ *R*^*D*^ is the D-dimensional feature vector of an example (precursor feature in MS1) and *y*_*i*_ ∈ {1 … *K*} is the corresponding class label (peptide sequence). For a query point **x** and a given distance metric *d*, a distribution over classes is computed by a softmax over distances to prototypes in the embedding space:

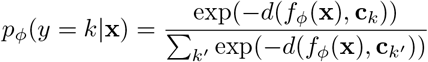

Prototypical Networks learn the parameters *ϕ*, of the mapping *f*_*ϕ*_ into the *M* -dimensional embedding space, by minimising the negative log-probability:

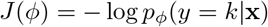

That is, the parameters of the mapping function are learnt such that the distance between the *M* -dimensional representations of data points from the same class and the mean vector of their embedding, **c**_*k*_, is minimized.

However, we observed that the learned embedding contains information on gradient length differences when two datasets were combined at training using the conventional protonet loss (Figure S1) - that is identifications belonging to the same class (modified sequence/charge) that exist in both datasets are not close on the embedding. Instead, we observed that embedding vectors of the identifications from the same study were very close to each other on the embedding space (Figure S1a). We addressed the dependence between the representations of data points from the same study by replacing the loss *J* (*ϕ*) with a conditional loss.

### The peptideprotonet model

We replace the loss *J* (*ϕ*) with *J* (*ϕ*|*z*):

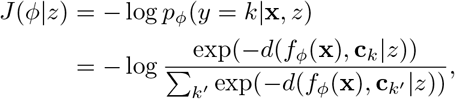

where *d*(*x, x*^∗^ | *z*) denotes the conditional distance; that is the Euclidean distance between two embedding vectors *x* and *x*^∗^, conditional on *z*, where *z* represents the study.

The conditional loss, which is inspired by conditional Variational Autoencoders (cVAEs) (Sohn et al., 2015), breaks the dependence between the representations of identifications from the same study and encourages instances from the same peptide class across the studies to be embedded more closely in the embedding space. In our particular case, we augment the embedding learned by *f*_*ϕ*_ with a categorical encoding of the study from which the support and query points are sampled at each episode. The conditional loss encourages *ϕ* to learn features that can reliably predict the same peptide identification in both short and long gradient data acquisitions.

Peptideprotonet is a supervised classification model. The embedding network consists of two linear layers (64 units and 10 units, respectively), with a single ReLU activation between layers, and is optimized with the Adam optimizer. The model were trained in 1-shot, 5-query, and 164-ways. At the validation and test steps the model operated in 1 test-shot, 5 test-query and 5 test-ways. That is - every time the model was presented with 1 support instance and 5 query instances from 164 peptide sequences at the training. At the test and validation time, the performance of the model was assessed in correctly embedding 1 support instance and 5 query instances from 5 peptide sequences. The model was trained for 300 epochs. For the few-shot classification, we randomly selected 90K labels (peptide sequence-charge) in the dataset for training and 20K for the validation dataset.

### Peptide Identity propagation in the embedding space

Let *v* = *f*_*ϕ*_ (**x**) denote the M-dimensional representation of query peptide **x**. We propagate peptide sequences from identified peptides to query peptides in each individual run based on the Euclidean distance of the representation *f*_*ϕ*_(**x**) of the query peptide **x** to a set of peptide prototypes, and agreement of the charges of query peptide and the proto-type - that is the closest prototype to *f*_*ϕ*_(**x**), weighted by the charge agreement between query and prototype. Let *N*_*v*_ be the set of peptide prototypes within a (*k*-nearest neighbors) neighborhood of *v*, and *c* denote a prototype in neighbor-hood *N*_*v*_. Let *n* denote the number of prototypes in *N*_*v*_. We compute standard deviation of distances between query embedding vector and the prototypes:

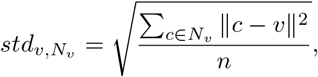

We then apply a Gaussian Kernel to the Euclidean distances:

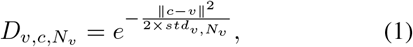

Let *Y* be the peptide label (i.e. peptide sequence/charge/modification). Let *η* denote the charge state. The probability that a query peptide belongs to peptide sequence *k*, given that the prototype *c*_*k*_ is in the neighborhood *N*_*v*_ and the embedding vector is determined as:

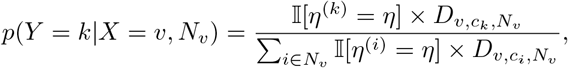

where *η*^(*i*)^ is the charge state of the *i*th prototype in the neighborhood, and *η* is the charge state of the query peptide. This ensures that the prototype selected for label propagation has the same charge state as the query peptide.

The peptide sequence with highest probability is assigned as the sequence of the query peptide:

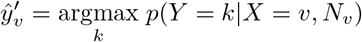

The uncertainty in the assignment is estimated using:

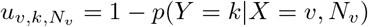

finally, if uncertainty is larger than a user-specific value, *κ*, the query precursor feature is considered as out-of-distribution (that is, the precursor feature does not belong to any of the identifications) and is discarded:

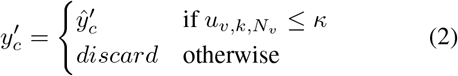

While the uncertainty score, 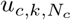, and the confidence score, *p*(*Y* = *k*|*X* = *c, N*_*c*_), are complementary, the earlier is useful for statistical modeling, whereas the latter is useful to express how certain we are about a transferred identification. For example, the uncertainty score can be used as precision weights in *limma*’s linear models to improve Empirical Bayes moderated t-statistics and differential abundance testing results. In this work, we mostly use the term confidence score to reflect the confidence in identification transfer.

### Two-pass operation to increase depth of coverage

The PIP framework can be extended beyond propagation of identifications between runs in a query dataset (match between run): to increase the depth of coverage. This is achieved by co-embedding the runs in a query dataset with one or more experimental libraries, similar to Data Independent Acquisitions (DIA), where experimental or predicted libraries are used to maximise identifications detected in all runs. Our propagation workflow operates in two-passes: in the first pass, if the confidence score for a precursor feature associated with a peptide sequence from the experiment is larger than 1 − *κ* threshold, the assignment is approved. If the confidence score is less than the threshold, the (embedded representation of) precursor feature is compared against (embedded representations of) identifications from the co-embedded libraries in the second pass. If the confidence score for an assignment from the co-embedded library was within the 1 − *κ* threshold, the peptide is transferred from the library to the query run. Peptides from new proteins that are not originally detected in the experiment can be transferred to runs by this approach, resulting in increase in depth of coverage. Note that in the paper, we have used *thr* to refer to the *κ* parameter for simplicity.

### Datasets

#### Model training using whole-proteome digests of five organisms and HeLa fractions acquired on diverse gradient lengths

We used 312 runs containing whole-proteome digests from five species (HeLa, Yeast, CElegans, Drosophila and Ecoli) published by (Meier et al., 2021) and 112 HeLa cell lysate and HeLa fractions runs published by (Meier et al., 2018) to train our model. The datasets are both acquired on timsTOF Pro mass spectrometer. (Meier et al., 2021) used a number of different enzymes to generate peptide fractions. They have used various gradient lengths of 30 min, 60 min and 120 min for data acquisition. In (Meier et al., 2018) data is acquired in 120 min gradient lengths. Details of sample preparation and data generation can be found in the original publications. We used the *evidence.txt* files generated by MaxQuant that were published by the authors to train our models (The ProteomeTools results were left out from Meier et al. data). Contaminants and Reverse Complement identifications, if any, were discarded. Charge one identifications, if any, were discarded. We only retained the feature with highest intensity, if multiple ions were reported for a peptide. Since our model training framework (1-shot, 5-query) required at least six training examples for any given peptide, peptides with six or more instances across the two datasets were retained in the training data. This resulted in a collection of 230,060 peptides and 34,411 proteins, available for training. We selected 90,000 peptide precursors at random for the training set, and 20,000 for validation. There was no intersection between peptides in training and validation splits. The following attributes were selected from the published *evidence* tables: `Charge, Mass, m/z, Retention time, Retention length, Ion mobility index, Ion mobility length` and `Number of isotopic peaks`. These attributes were scaled to have mean zero and unit standard deviation prior to model training. Our model learns a 10-dimensional representation of these attributes.

#### Assessment of quantification precision using four HeLa cell lysate replicates

We used the four HeLa cell lysate replicates published by Meier et al to evaluate precision and sensitivity of peptide and protein abundances in PIP results. The runs were acquired on a timsTOF Pro mass spectrometer with 100 ms accumulation time. Details of sample preparation and data generation can be found in the original publication. We used the published *allPeptides.txt* and *evidence.txt* results for PIP analysis (PRIDE accession PXD010012). We only retained identifications detected by MULTI-MSMS in MaxQuant results. The *evidence* tables from MaxQuant and PIP results were then processed as follow: We only retained the feature with highest intensity, if multiple ions were reported for a peptide in the *evidence* table. Contaminants and Reverse Complement identifications, if any, were discarded. Charge one identifications, if any, were discarded. Peptide intensities were log_2_ transformed and normalized by Quantile Normalization implemented in limma. Protein abundance was calculated by Top-N approach, that is we sum the three highest intensity peptides to quantify the abundance of the protein. We used this dataset to assess quantification CV at the peptide- and protein-level, and to examine improvements in data completeness.

#### Evaluation of quantification precision and false transfer rate using ten HeLa cell lysate replicates

We used the ten HeLa cell lysate replicates published by Prianichnikov et al. (2020) to evaluate peptide precision and FDR in PIP results. The runs were acquired on a timsTOF Pro mass spectrometer on a 2hr gradient. Details of sample preparation and data generation can be found in the original publication. We used the published *allPeptides.txt* and *evidence.txt* results for PIP analysis (PRIDE accession PXD014777). We only retained identifications detected by MULTI-MSMS in MaxQuant results. The *evidence* tables from MaxQuant and PIP results were then processed as follow: We only retained the feature with highest intensity, if multiple ions were reported for a peptide in the *evidence* table. Contaminants and Reverse Complement identifications, if any, were discarded. Charge one identifications, if any, were discarded. Peptide intensities were log_2_ transformed and normalized by Quantile Normalization implemented in limma. Protein abundance was calculated by Top-N approach, that is we sum the three highest intensity peptides to quantify the abundance of the protein.

#### Assessment of false transfers in biologically heterogeneous dataset of T cells secretome OSCC

This dataset contains six runs comparing secretome from tumor microenvironment of Oral squamous cell carcinoma (OSCC) (n=3) to non-malignant samples (n=3). The study specifically investigated the T cells population in secretome, hence, it is referred to as T cells OSCC dataset in this work. The runs were acquired on a timsTOF Pro mass spectrometer. Raw spectral .d files were downloaded from accession PXD023049 from PRIDE repository, searched and quantified by MaxQuant version 1.6.6.0. as per procedures described by the authors on the PRIDE project repository. Details of sample preparation and data generation can also be found on PRIDE. We used the *allPeptides.txt* and *evidence.txt* for PIP analysis. The *evidence* tables from MaxQuant and PIP results were then processed as follow: We only retained the feature with highest intensity if multiple ions were reported for a peptide in the *evidence* table. For the heatmap, peptide intensities were log_2_ transformed and retained in the study if they were not missing in more than two samples. Also, the modifications were excluded from the heatmap analyses. We used this dataset to assess performance in the presence of biological variability in small experiments, when peptides could be missing not at random.

#### Evaluation of quantification accuracy using a three-organism hybrid proteome dataset

We used a three-organism proteome dataset to compare quantification accuracy in MaxQuant MBR+ and PIP results. This dataset is published by Prianichnikov et al, and contains six runs from two conditions A and B, in which Human, Yeast and E.coli proteins are spiked at known ratios. The expected ratios in B versus A are 1:1 (Human), 1:2 (Yeast) and 4:1 (E.coli) proteins. The data were acquired on a timsTOF Pro mass spectrometer. Details of sample preparation and data generation can be found in the original publication. We used MaxQuant results published by them with MBR enabled. The PIP analysis was done using the published *allPeptides.txt* and *evidence.txt* results (PRIDE accession PXD014777). We only retained identifications detected by MULTI-MSMS in MaxQuant MBR+ results. We also tried the published MaxQuant MBR-results, but since there was no difference between the two results, the MBR+ results were used for consistency. The *evidence* table from MaxQuant MBR+ and PIP results were then processed as follow: We only retained the feature with highest intensity, if multiple ions were reported for a peptide in the *evidence* table. Contaminants and Reverse Complement identifications were discarded. Charge one identifications, if any, were discarded. Protein abundance was calculated by Top-N approach, that is we sum the three highest intensity peptides to quantify the abundance of the protein. Protein abundances were log_2_ transformed and normalized by Quantile Normalization implemented in limma. Proteins ending with_YEAST in the protein name were classified as Yeast proteins. Proteins with a ECOLI ending in their names were classified as E.coli, and the rest of the unannotated proteins were classified as HUMAN proteins, as per original publication. A pair of samples from conditions A and B were randomly selected to estimate peptide fold-changes in B vs. A. Average abundance was computed by averaging log_2_ abundances across all runs.

## Acknowledgements

We thank Rune Larsen and Daryl Wilding-McBride from WEHI proteomics for helpful discussions.

S.H-z is supported by an Australian Postgraduate Award.

## Data availability

All of the datasets analyzed in this manuscript are public and published in other papers. We have referenced them in the manuscript. The model and code to reproduce the results of this paper is also available at https://github.com/DavisLaboratory/peptideprotonet_reproducibility.

## Author contributions

S.H-z conceived and designed the framework under supervision of M.J.D and A.I.W. S.H-z and J.M wrote the code. SH-z performed the analyses. All authors wrote and approved the manuscript.

## Competing interests

There are no competing interests to disclose.

## Notes

### Competing Interest Statement

The authors have declared no competing interest.

